# Compensating for intersegmental dynamics across the shoulder, elbow and wrist joints during feedforward and feedback control

**DOI:** 10.1101/115675

**Authors:** Rodrigo S. Maeda, Tyler Cluff, Paul L. Gribble, J. Andrew Pruszynski

## Abstract

Moving the arm is complicated by mechanical interactions that arise between limb segments. Such intersegmental dynamics cause torques applied at one joint to produce movement at multiple joints and, in turn, the only way to create single joint movement is by applying torques at multiple joints. Here, we investigated whether the nervous system accounts for intersegmental limb dynamics across the shoulder, elbow, and wrist joints during self-initiated reaching and when countering external mechanical perturbations. Our first experiment tested whether the timing and amplitude of shoulder muscle activity accounts for interaction torques produced during single-joint elbow movements from different elbow initial orientations and over a range of movement speeds. We found that shoulder muscle activity reliably preceded movement onset and elbow agonist activity, and was scaled to compensate for the magnitude of interaction torques arising because of forearm rotation. Our second experiment tested whether elbow muscles compensate for interaction torques introduced by single-joint wrist movements. We found that elbow muscle activity preceded movement onset and wrist agonist muscle activity, and thus the nervous system predicted interaction torques arising because of hand rotation. Our third and fourth experiments tested whether shoulder muscles compensate for interaction torques introduced by different hand orientations during self-initiated elbow movements and when countering mechanical perturbations that caused pure elbow motion. We found that the nervous system predicted the amplitude and direction of interaction torques, appropriately scaling the amplitude of shoulder muscle activity during self-initiated elbow movements and rapid feedback control. Taken together, our results demonstrate that the nervous system robustly accounts for intersegmental dynamics, and that the process is similar across the proximal to distal musculature of the arm as well as between feedforward (i.e., self-initiated) and feedback (i.e., reflexive) control.

**NEW & NOTEWORTHY:** Intersegmental dynamics complicate the mapping between applied joint torques and the resulting joint motions. Here, we provide evidence that the nervous system robustly predicts these intersegmental limb dynamics across the shoulder, elbow and wrist joints during reaching and when countering external perturbations.

## INTRODUCTION

Most arm movements require the nervous system to coordinate multiple joints. Complicating this coordination are mechanical interactions between limb segments that arise because torques generated at one joint cause rotational forces at other joints, and thus produce motions without muscle contraction (Hollerbach and Flash, 1982). For a two-link arm in the horizontal plane, for example, applying torque only at the elbow will cause both the shoulder and elbow to move. Thus, the only way to produce a single joint elbow movement is to generate torque at both the shoulder and elbow joints.

Many researchers have examined how the nervous system accounts for the arm’s intersegmental dynamics during self-initiated (i.e. voluntary) reaching movements (Almeida et al., 1995; Cooke and Virji-Babul, 1995; Corcos et al., 1989; Galloway and Koshland, 2002; Gottlieb, 1998; Gribble and Ostry, 1999; Gritsenko et al., 2011; Hollerbach and Flash, 1982; Koshland et al., 1991; Pigeon et al., 2013; Sainburg et al., 1995, 1999; Virji-Babul and Cooke, 1995). As a whole, these studies clearly indicate that control signals sent to arm muscles appropriately predict upcoming interaction torques, and thus, likely rely on an internal model of mechanical interactions between limb segments (Wolpert and Flanagan, 2001). For example, in the context of both constrained and unconstrained single-joint movements, Almeida et al. (1995) demonstrated that muscle activation patterns at a stationary joint (either the shoulder or elbow) counteract the interaction torques that arise because of the motion of the adjacent joint, and thus prevent their movement. Gribble and Ostry (1999) further showed that shoulder muscle activity predictively accounts for interaction torques by showing that shoulder muscle activity occurs prior to movement onset and scales appropriately with differences in the speed and amplitude of single joint elbow movements.

Several groups have also investigated whether rapid feedback responses (i.e. reflexes) are modulated in a way that accounts for intersegmental dynamics (Crevecoeur et al., 2012; Kurtzer et al., 2008, 2009, 2014, 2016; Lacquaniti and Soechting, 1984, 1986a, b; Pruszynski et al., 2011; Soechting and Lacquaniti, 1988). For example, Kurtzer and colleagues 2008 applied a combination of shoulder and elbow torque perturbations that led to minimal shoulder motion but different amounts of elbow motions. This approach allowed them to test whether rapid feedback responses in shoulder muscles were dependent on local stretch information or whether they accounted for the limb’s dynamics and thus responded to the applied torques. Their results showed that the short-latency feedback response (20-50ms post-perturbation), which is mediated by spinal circuits, responded only to local joint motion. In contrast, the long-latency feedback response (50-100ms post-perturbation), which is partially mediated by the same cortical structures that contribute to voluntary control (Pruszynski and Scott, 2012; Shemmell et al., 2010), responded to the underlying applied torques. This is consistent with the idea that long latency responses are organized based on an internal model of intersegmental dynamics.

Although these previous studies are consistent with the idea that both voluntary and feedback control of the arm make use of an internal model of the limb’s dynamics, they have focused their investigations on a limited set of parameters in the context of two joints - usually the shoulder and elbow. Here, we performed four experiments with a three degree-of-freedom exoskeleton robot that allows flexion and extension movements of the shoulder, elbow, and wrist joints. Our aim was to examine how the nervous system accounts for intersegmental dynamics across these three joints during voluntary reaching movements and when responding to unexpected mechanical perturbations. In our first experiment, we extended the work of Gribble and Ostry (1999) and tested whether shoulder muscles compensate for the magnitude of interaction torques arising during single-joint elbow movements performed across a range of speeds from different initial elbow orientations. In our second experiment, we tested whether elbow and shoulder muscles compensate for interaction torques introduced by single-joint wrist movements. In our third and fourth experiments, we used the same paradigm to investigate whether voluntary and rapid feedback control of shoulder muscles compensates for interaction torques introduced by changing the orientation of the hand. Taken together, we found that both voluntary and feedback control of upper limb muscles predictively compensates for interaction torques arising across the shoulder, elbow and wrist joints.

## MATERIALS AND METHODS

### Participants

A total of sixty participants (aged 18-38; 32 males, 28 females) with no known musculoskeletal or neurological diseases participated in the studies described here. Each participant was tested in one of the four experiments. All participants had normal or corrected-to-normal vision and self-reported that they were right hand dominant. The Office of Research Ethics at Western University approved all experimental procedures according to the Declaration of Helsinki, and all participants gave informed written consent prior to participating in an experiment.

### Apparatus

Experiments were performed with a three degree-of-freedom exoskeleton robot (Interactive Motion Technologies, Boston, MA) that allows flexion/extension rotations of the shoulder, elbow, and wrist joints in a horizontal plane intersecting the shoulder joint (for details, see Weiler et al., 2015, 2016). The exoskeleton measures movement kinematics of the shoulder, elbow and wrist joints, and can apply torques that flex or extend each independent joint. Visual stimuli were projected downward from a 46-inch LCD monitor (60 Hz, 1,920 - 1,080 pixels, Dynex DX-46L262A12, Richfield, MN) onto a semi silvered mirror. The mirror was mounted parallel to the plane of motion and blocked direct vision of the participant’s arm. Visual feedback about the participant’s hand position was provided in real time by displaying a cursor (purple circle, 1 cm diameter) at the location of the exoskeleton handle on the visual display. Each segment length of the robot was adjusted to fit the participant’s arm, so the cursor was aligned with their actual hand position. The lights in the experimental room were extinguished for the duration of data collection.

### Experiment 1: Compensating for interaction torques during single joint elbow movements

Fifteen participants performed thirty degrees of elbow flexion and extension movements, starting from three different initial elbow orientations, and at two movement speeds. Each trial began with the participant grasping the exoskeleton handle and moving the projected cursor to a blue circle (i.e., home target: 2 cm diameter) that corresponded to the hand position when the shoulder and elbow joints were at a specific orientation for each condition. We displayed the home target so the shoulder joint was always positioned at 60°(external angle) and the initial orientation of the elbow could be positioned at either 45°, 60°or 75°(external angles) for elbow flexion movements and 75°, 90°, or 105°angles for elbow extension movements (Figure 1, top left panel). The wrist joint was physically locked at 16°in all conditions, including flexion and extension movements. Once participants achieved and remained at one of these joint orientations for 3 s, an instruction about movement speed (“fast”, or “slow”) was displayed 2 cm above the home target. At the same time, the hand feedback cursor was extinguished and remained off for the rest of the trial. After a short delay (0-1 s, uniform distribution), a white goal target was drawn such that it could be reached with pure elbow flexion or extension (30°). The participant’s task was to move the hand into the goal target at the instructed speed. The goal target turned green when movement time, calculated as the time from exiting the start position to entering the goal position, was less than 120ms and 220ms for fast and slow movements, respectively. If movement speed exceeded 220ms, the goal target turned red. The order of all elbow orientations and movement speeds were randomized. Participants completed 480 trials total (2 speeds × 2 directions × 3 initial elbow configurations × 40 repeats per condition). About 2.5h was required to complete Experiment 1.

**Figure 1:**
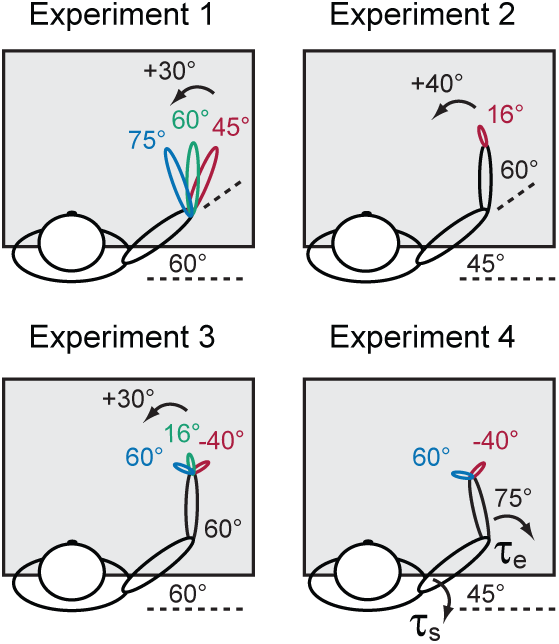
Experimental setup. Participants were seated and placed their right arm in a three-joint (shoulder, elbow, wrist) robotic exoskeleton that allowed them to perform reaching movements in the horizontal plane. A semi-silvered mirror occluded vision of their hand. For clarity, only flexion movement conditions for each experiment are shown, as depicted by the arrows in Experiments 1, 2 and 3. Arrows in Experiment 4 represent the multi-joint step-torques applied to the shoulder and elbow joints.

### Experiment 2: Compensating for interaction torques during single joint wrist movements

Fifteen participants performed 40°wrist flexion and extension movements. In this experiment, participants placed their hand into a custom made hand trough attached to the exoskeleton handle, which kept their fingers straight. Each trial began with the participant moving their hand to a blue circle (i.e., home target: 2 cm diameter) that corresponded to the tip of their index finger when the shoulder, elbow and wrist joints were at 45°, 60°and 16°angles, respectively (Figure 1). Once participants held this posture for 3 s, the cursor indicating the position of the tip of the index finger was extinguished and a white goal target was drawn after a short random delay (0-1 s, uniform distribution), such that it could be reached with 40°of wrist flexion alone (Figure 1, top right panel). We instructed participants to move the cursor from the home target into the goal target. The goal target turned green when the movement between these targets was less than 150 ms and turned red otherwise. Feedback of the index fingertip position remained off during the movement and re-appeared to indicate the start of a new trial. The previous goal target then became the new starting location for a wrist extension movement and the same sequence of events took place. Participants completed a total of 120 trials (60 wrist flexion and 60 wrist extension). About 1 h was required to complete the experiment.

### Experiment 3: Compensating for interaction torques introduced by hand orientation

Fifteen participants performed 30°elbow flexion and extension movements with different hand configurations. Participants used the custom-made hand trough as in Experiment 2. Participants moved their hand to a blue circle (i.e., home target: 2-cm diameter) that corresponded to the tip of their index finger when the shoulder and elbow were at 60° and the wrist was at −30°, 16°or 50°configurations (Figure 1, bottom left panel). Once participants achieved and remained at one of these joint configurations for 3 s, feedback of index fingertip position was extinguished and a white goal target was then drawn such that it could be reached with elbow rotation alone. The participant’s task was to move the hand between the targets within 150 ms while keeping the hand at the specified configuration. The goal target turned green when movement time was within the required time, and turned red when it was slower. The order of all wrist configurations was randomized. Participants completed 360 trials total (2 directions × 3 wrist configurations × 60 repeats per condition). About 2.5 h was required to complete Experiment 3.

### Experiment 4: Rapid feedback responses at the shoulder account for interaction torques caused by hand orientation

Fifteen participants were instructed to respond to multi-joint mechanical perturbations that led to similar shoulder and elbow motion profiles (see Kurtzer et al., 2008) while their hand was positioned in one of two initial orientations. All participants used the hand trough as in Experiments 2 and 3. At the beginning of a trial, participants moved the tip of the index finger (indicated by a cursor) to a blue circle (i.e., start target: 4 cm diameter) whose center was located above the center of rotation of the participant’s wrist when the shoulder and elbow were at 70°and 90°, respectively. After entering the start target, the exoskeleton gradually applied (over 2 s) a background torque of −2 Nm to the elbow and locked the wrist into two distinct configurations (-40°and 60°). A blue circle was then displayed (i.e., home target: 2 cm diameter) centered at the tip of the participant’s index finger when the shoulder and elbow were at 45°and 75°, respectively. Participants were instructed to move the cursor to the home target while counteracting the background load at the elbow.

After maintaining the cursor in the home target for a randomized duration (1.0-2.5 s, uniform distribution), the cursor was removed, and a step-torque (i.e., perturbation) was applied to the shoulder and elbow joints, which rapidly displaced the participant’s hand outside the home target. Critically, we applied a specific combination of torques at the shoulder and elbow so that this perturbation would lead to similar motion profiles at the shoulder and elbow joints in the two wrist configurations (Figure 1, bottom right panel). The torques that generated these motion profiles (minimal shoulder motion and substantial elbow motion) were −0.5 Nm and −5.5 Nm torques at the shoulder and elbow when the wrist was locked at −40°and −0.3 Nm and −5.5 Nm torques at the shoulder and elbow when the wrist was locked at −60°, respectively (i.e. Load Combination 1). We also coupled each of these torques with the other wrist configuration (i.e. Load Combination 2). Note that trials from Load Combination 2 were only present to ensure the perturbation was unpredictable and were not analyzed in detail. After perturbation onset participants were instructed to quickly bring the cursor back into a goal target (4 cm diameter centered on the home target). If the participant moved the cursor into the goal target within 375 ms of perturbation onset, the target circle changed from white to green, otherwise the target circle changed from white to red. Regardless of trial outcome, all torques were gradually removed 1300 ms after perturbation onset. The order of all wrist configurations and perturbations was randomized. Participants completed 300 trials total (2 torques × 2 wrist configurations × 75 repeats per condition). About 2.5 h was required to complete Experiment 4.

### Common experimental features

Before data collection for all experiments, participants performed normalization trials. In these trials, participants were instructed to move the cursor to a blue circle (i.e., home target: 2 cm diameter) that was centered at the robot’s handle (Experiment 1) or tip of the participant’s index finger (Experiments 2-4) when the shoulder, elbow and wrist joints were at 45°, 60°and 16°, respectively. Once in the home target, the exoskeleton gradually applied torques to either the shoulder, elbow or wrist joints, which plateaued at a constant ±2 Nm torque. Participants were instructed to counter these torques while maintaining the cursor in the home target for 4 s. Following this period, the joint torques were turned off. The order of the normalization trials, which included flexion and extension at each of the three joints, was randomized. Participants completed 4 trials of each condition.

For all experiments, rest breaks were given throughout or when requested. Prior to data collection participants completed practice trials until they comfortably achieved ~70% success rates (approx. 10 min).

### Muscle activity

In all experiments we collected muscle activity using surface EMG electrodes (Delsys Bagnoli-8 system with DE-2.1 sensors, Boston, MA). The participants’ skin was abraded with rubbing alcohol, and contacts were coated with conductive gel. Electrodes were then placed on the skin surface overlying the belly of six muscles for Experiments 1, 2 and 3 (pectoralis major clavicular head, PEC, shoulder flexor; posterior deltoid, DELT, shoulder extensor; biceps brachii long head, BI, shoulder and elbow flexor, wrist supinator; triceps brachii lateral head, TRI, elbow extensor; flexor carpi ulnaris, WF, wrist flexor; extensor carpi radialis, WE, wrist extensor). Electrodes were oriented parallel to the orientation of muscle fibers. All but wrist muscles were also recorded in Experiment 4. A reference electrode was placed on the participant’s left clavicle. EMG signals were amplified (gain = 103), band-pass filtered (20-450 Hz), and then digitally sampled at 2,000 Hz. Normalization trials (see above) prior to each experiment were used to normalize muscle activity such that a value of 1 represents a given muscle sample’s mean activity when countering a 2 Nm torque (see Pruszynski et al., 2008).

### Data analysis

All joint kinematics (i.e. hand position, and joint angles) were sampled at 500 Hz and then low-pass filtered (12 Hz, 2-pass, 4th-order Butterworth). EMG data were band-pass filtered (20-500 Hz, 2-pass, 2nd-order Butterworth) and full-wave rectified. For scoring the onset of phasic EMG bursts, the rectified signals were low-pass filtered (50 Hz, 2-pass, 12th-order Butterworth). For Experiments 1, 2, and 3 all data were aligned on movement onset. Movement onset was defined as 5% of peak angular velocity of the elbow joint for Experiment 1 and 3, and 5% of peak angular velocity of the wrist joint for Experiment 2. All data were aligned on perturbation onset for Experiment 4.

In Experiments 1, 2 and 3, we assessed whether muscles at stationary joints compensate for interaction torques as a function of limb configuration and speed. To compare the amplitude of muscle activity across these different conditions, we calculated the mean amplitude of phasic muscle activity across a fixed time-window (see Debicki and Gribble, 2005). We used a fixed time window of −200 ms to +100 ms relative to movement onset in Experiments 1 and 3, and −100 ms to +100 ms relative to movement onset in Experiment 2. These windows were chosen to capture the agonist burst of EMG activity in each of the experiments and our results did not qualitatively change with small changes in the start and end of the analysis window.

We assessed the timing of muscle activity by calculating the onset of the first phasic EMG burst of each muscle in each trial. For each trial we computed baseline EMG activity over a fixed 100-ms window, 400 ms before the start of movement. The onset of the first EMG burst was scored as the time at which the EMG signal rose three standard deviations above the mean baseline level and remained above that level for 50 ms. A research assistant, unaware of the study hypotheses, performed additional manual inspection of the results of this algorithm and confirmed, rejected (10%) or remarked these onsets values relative to movement onset.

In Experiment 4 we investigated whether shoulder muscles compensated for limb dynamics related to hand orientation when countering mechanical perturbations. To test whether the short and long-latency stretch response of a shoulder flexor muscle accounts for hand orientations, we binned PEC EMG into previously defined epochs of time (Pruszynski et al., 2008). This included a pre-perturbation epoch (PRE, −50-0 ms relative to perturbation onset), the short-latency stretch response (R1, 25-50 ms), the long-latency stretch response (R2/3, 50-100 ms), and the so-called voluntary response (VOL, 100-150 ms).

Data processing was performed using MATLAB (The Mathworks, Natick, MA) and statistical analyses were performed using R (RStudio, Boston, MA). We performed different statistical tests (e.g., repeated measures ANOVA, paired and single-sample t-tests, and Tukey tests for multiple comparisons) when appropriate for each of the four experiments. Details of these procedures are provided in the Results. Experimental results were considered statistically significant if *p <* 0.05.

## RESULTS

### Experiment 1: Compensating for interaction torques during single joint elbow movements

We instructed participants to move the tip of their index finger between two targets. The start and end targets were always positioned so that flexing or extending only the elbow joint would successfully transport the hand between the two targets (Figure 1). We manipulated the start and end target location, as well as the required movement speed, to examine whether shoulder muscle activity compensates for variations in the amplitude of interaction torques that arise as a function of the elbow’s initial configuration and the speed of rotation. Participants quickly learned the task and had little difficulty reaching the goal target within the imposed speed and accuracy constraints (mean success rate = 90%). We included all trials in the analysis. Although we did not explicitly enforce a particular trajectory between the two targets, participants achieved the required movement by rotating their elbow joint while keeping the shoulder and wrist joints relatively fixed (Figure 2 A,B).

**Figure 2:**
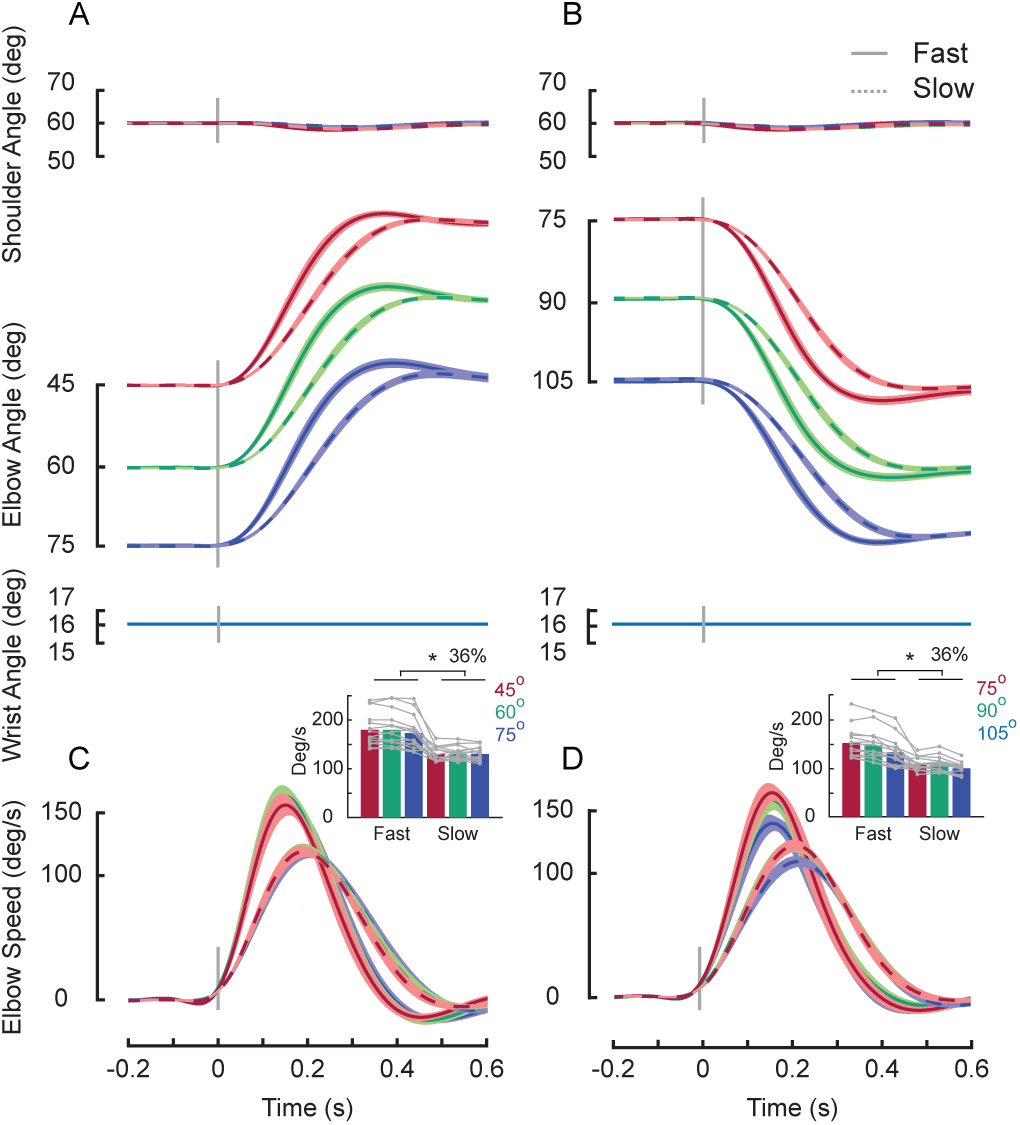
Reaching Behavior, Experiment 1. (A) Kinematics of single joint elbow movements is shown for three different elbow configurations and two movement speeds. Thick lines represent the group mean of the kinematics of the shoulder, elbow and wrist joints during elbow flexion. Fast and slow movements are represented as solid and dashed lines, respectively. Shaded areas represent the standard error of the mean. Data are aligned on movement onset. (B) Kinematics of extension movements are shown using the same format in A. (C) Joint velocities are shown. The inset summarizes peak velocity as a function of elbow configuration and instructed movement speed. Bars represent the mean peak velocity across participants and each gray line represents the mean peak velocity of a single participant. The asterisk indicates a reliable effect of speed (*p <* 0.05, see main text). (D) Extension movements are shown.

We used two-way repeated-measures ANOVAs (one for elbow flexion and one for elbow extension) to test whether the peak velocity of elbow rotation differed between our three elbow orientation and two speed conditions. For both elbow flexion (Figure 2C) and extension movements (Figure 2D), we found a reliable effect of speed condition on peak elbow rotation velocity (flexion: *F*_1,78_ *=* 23.7, *p* < 0.0001; extension: *F*_1,78_ *=* 25.6, *p* < 0.0001, but no significant effect of elbow orientation (*F*_2,78_ = 0.006, *p* = 0.99; *F_2_,_78_ =* 1.134, *p* = 0.32) or interaction (*F*_2_,_78_ = 0.053, *p =* 0.94; *F*_2_,_78_ = 0.044*,p =* 0.95). On average, fast movements were 36% faster than slow movements for both flexion and extension directions.

Figure 3 shows the group average shoulder flexor (PEC, panels A and B) and extensor (DP, panels D and E) muscle activity associated with fast (left column) and slow (right column) movements at each initial elbow configuration. Despite minimal shoulder rotation, we found substantial shoulder muscle activity prior to movement onset. We found no evidence of substantial co-contraction at the shoulder; rather, agonist shoulder muscle activation appeared at the muscle appropriate for counteracting the interaction torques that were about to arise due to the upcoming forearm rotation (i.e. shoulder flexors for elbow flexion and shoulder extensors for elbow extension). We quantified the magnitude of agonist shoulder muscle activity in each condition by computing the mean muscle activity over a fixed time window (-200 to 100 ms) relative to movement onset (see Methods). For each participant, we related the agonist muscle activity to the three initial joint configurations using linear regression (Figure 3C,D). We predicted a negative relationship between muscle activity and initial external angle as increasing the external angle decreases the interaction torques at the shoulder (Gribble and Ostry, 1999). Consistent with our prediction, a one-sample t-test of each individual’s slope revealed a reliable negative relationship for both fast and slow elbow flexion (fast: *t*_14_ = −6.5, *p* < 0.0001; slow: *t*_14_ = –4.361, *p* = 0.0006) and extension movements (*t*_14_ = –2.39, *p* = 0.03; *t*_14_ = –3.36, *p* = 0.004). We also predicted larger shoulder muscle activity for the faster movement speed as interaction torques at the shoulder scale with elbow rotation velocity. A one-sample t-test of each individual’s intercept confirmed our prediction in flexion and extension movements (flexion: *t*_14_ = 5.82, *p* < 0.0001; extension: *t*_14_ *=* 5.64,p < 0.0001).

**Figure 3:**
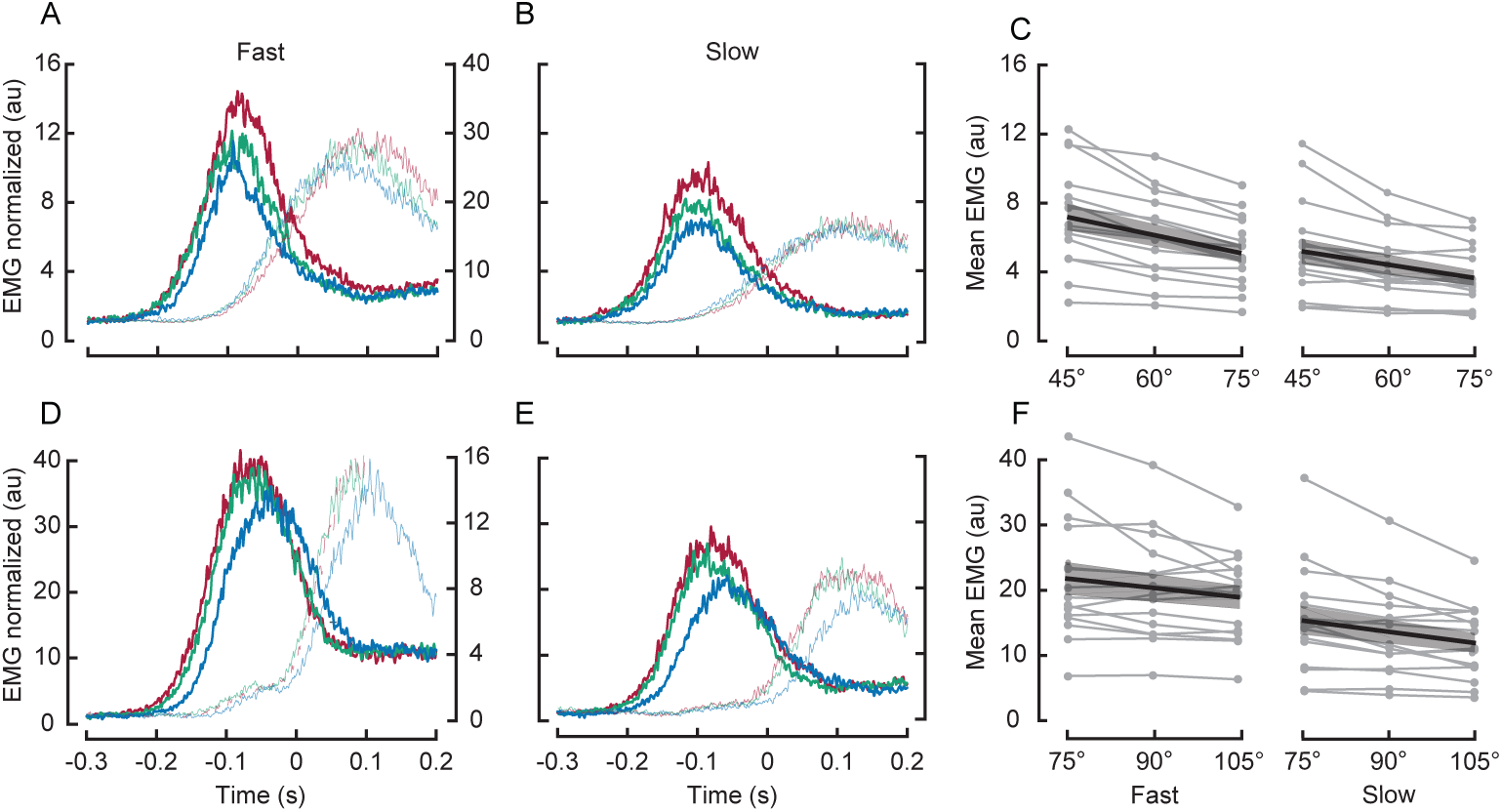
Muscle Activity, Experiment 1. (A) Thick and thin lines represent average agonist (PEC) and antagonist (PD) muscle activity during fast flexion movements. Initial configuration is depicted by line color. EMG data are normalized as described in the Methods. Data are aligned on movement onset. (B) EMG data are shown using the same format as A for movements in the slow condition. (C) Individual data (gray lines) and mean regression slopes (black line; shaded area = SE) between mean agonist muscle activity and elbow orientations during elbow flexion movements in different speeds. Mean agonist muscle is shown in a fixed time window (-200 ms to 100 ms relative to movement onset). (D, E, F) Elbow extension movements are shown using the same format as (A, B, C). Agonist = PD; antagonist = PEC, respectively.

Figure 4 shows a histogram of all shoulder and elbow muscle onset times for all subjects relative to movement onset for elbow flexion (A,B) and extension (D,E) trials. These histograms indicate that shoulder muscle activity almost always begins well before movement onset (i.e. elbow rotation). For elbow flexion and extension trials, PEC and PD muscle activity reliably preceded movement onset by 160 ms (SE 6) (*t*_14_ = –26.5, *p <* 0.0001; one-sample t-test versus 0) and 150 ms (standard error of the mean, SE 4) (*t*_14_ = –38.1, *p* < 0.0001), respectively. To examine differences in onset times between shoulder and elbow muscles, Figure 4C,F overlay the cumulative distribution function of shoulder and elbow muscle onset times relative to movement onset. Statistical analysis revealed that PEC onsets reliably preceded BB onsets by a modest 10 ms (SE 4; *t*_14_ = –2.4, *p* = 0.02) for flexion trials. On the other hand, we found no statistically reliable difference between onset times in PD and TB muscles for extension trials (*t*_14_ = –1.6, *p* = 0.14) though the data showed a similar trend.

**Figure 4:**
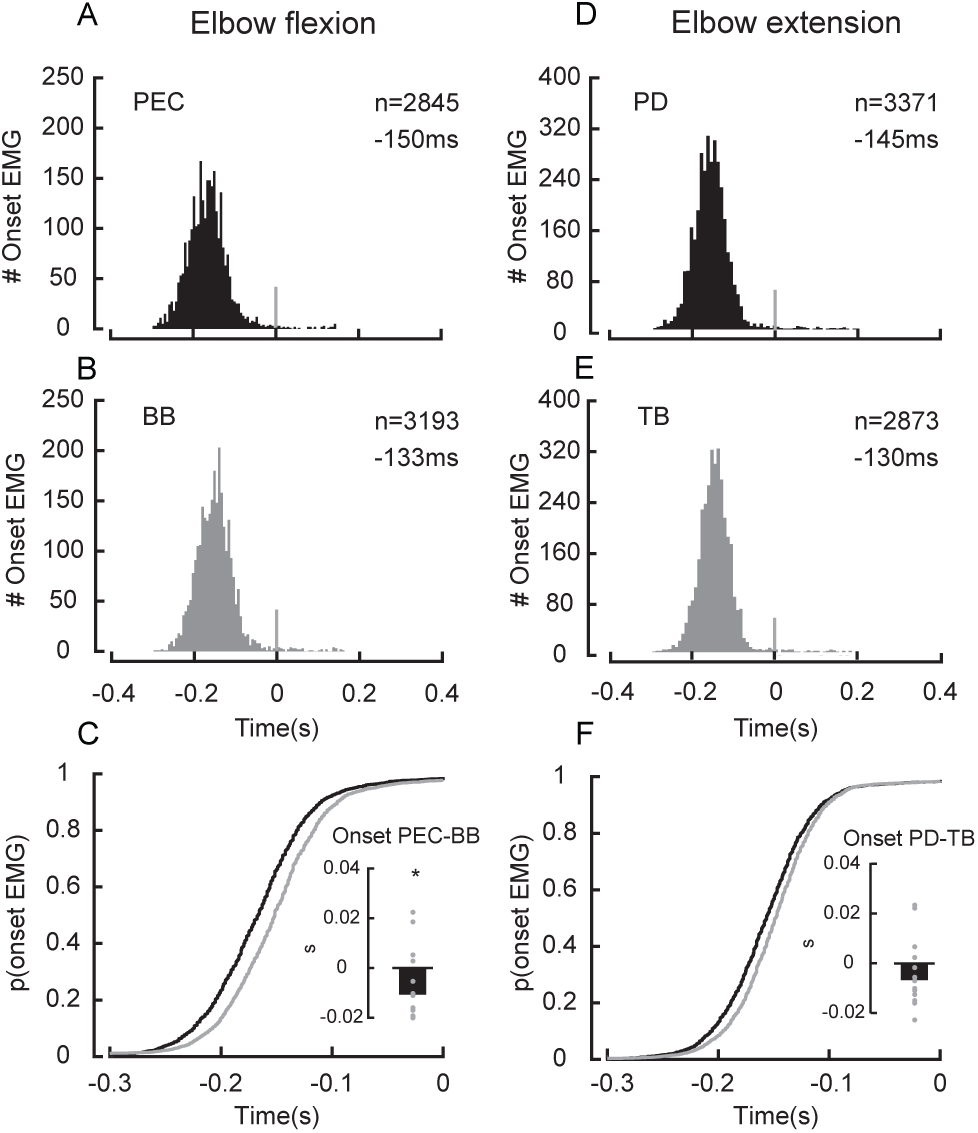
Muscle Onset Timing, Experiment 1. (A) Histogram of onset times of PEC muscles and (B) histogram of onset times of BB muscles during elbow flexion movements. Data are aligned on movement onset. (C) Cumulative distributions of onset times of shoulder and elbow muscles are shown for elbow flexion movements. The inset shows the difference between mean shoulder and elbow onset times. (D - F) elbow extension movements are shown using the same format as (A - C).

Lastly, we examined the correlation between shoulder and elbow muscles in terms of their amplitude of activation (Figure 5A,B) and onset time (Figure 5C,D) across movements. We predicted a positive relationship between shoulder and elbow muscle activity magnitude and onset timing, consistent with a process that actively compensates for intersegmental dynamics on a trial by trial basis. As predicted, in terms of magnitude, all 15 participants showed a significant positive correlation for flexion movements and 14 participants showed a significant positive correlation for extension movements. Similarly, in terms of timing, 14 of 15 participants showed a significant positive correlation for both flexion and extension movements.

**Figure 5:**
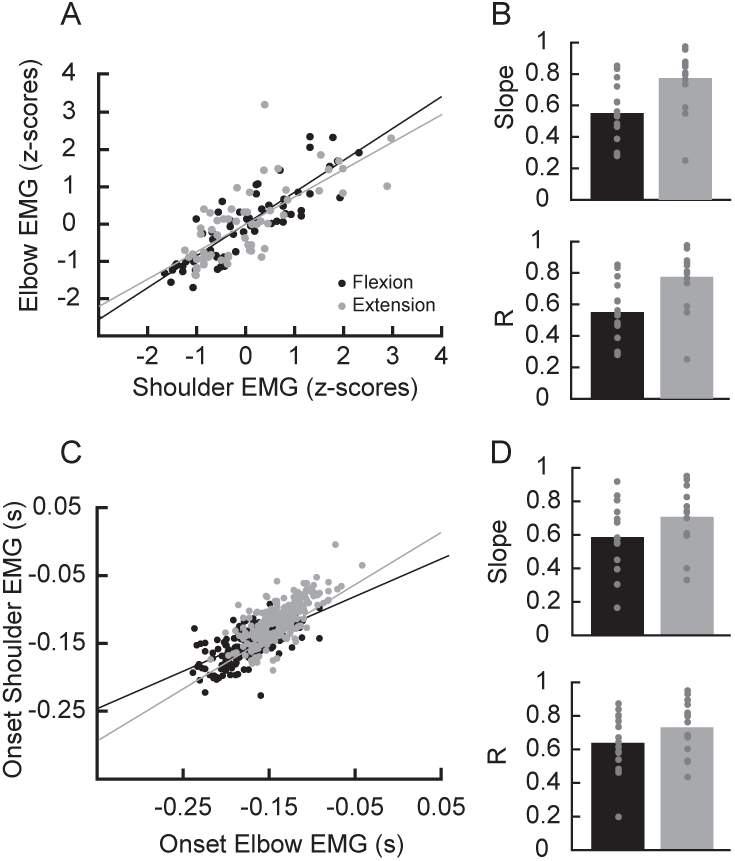
Trial-by trial relationships between muscles, Experiment 1. (A) Exemplar correlation between muscle activity at the shoulder and elbow. Each dot represents mean muscle activity in pre-defined epoch from a single trial (-200 to 100 ms relative to movement onset). Values are z-normalized. (B) Group mean slopes and correlation coefficients are shown. Each dot represents data from a single participant. (C and D) Muscle onset times are shown using the same layout as (A,B).

### Experiment 2: Compensating for interaction torques during single joint wrist movements

We investigated whether elbow muscles account for interaction torques introduced by wrist movements. The start and end targets were always placed along an arc centered on the wrist joint so that flexing or extending only the wrist joint would successfully transport the hand between the targets (Figure 1). Participants quickly learned the task and had little difficulty reaching the goal target within the imposed speed and accuracy constraints (mean success rate = 99%). Figure 6 shows the average kinematics of the elbow and wrist joints during single-joint wrist flexion and extension movements, respectively. Similar to Experiment 1, although we did not give explicit instructions about how the participants should complete the required movement, participants did so by almost exclusively rotating the wrist joint.

**Figure 6:**
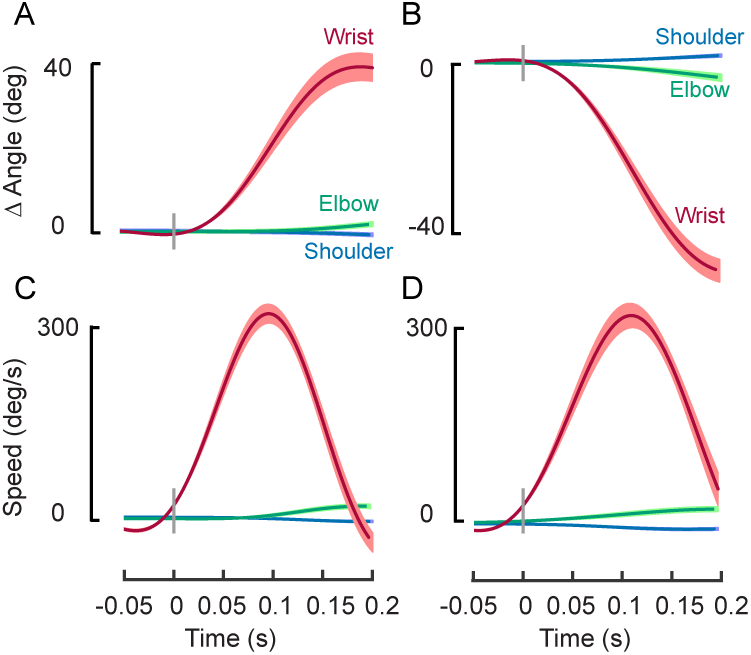
Reaching Behavior, Experiment 2. (A,B) Average kinematic profiles for the shoulder, elbow and wrist joints for flexion and extension conditions in Experiment 2, respectively. Data are aligned on movement onset. Shaded areas represent the standard error of the mean. (C and D) Joint velocities are shown using the same format as A and B.

Figure 7 shows average muscle activity for wrist flexion (A,B) and wrist extension movements (D,E). These plots indicate that elbow muscle activity preceded movement onset, and that these were directional responses appropriate for counteracting the interaction torques about to arise due to upcoming hand rotation. We quantified this compensation by computing the mean muscle activity in a fixed time window (-100 to 100ms) relative to movement onset. For wrist flexion movements (Figure 7C), a one-sample t-test showed that both the WF (*t* _14_ = 6.3, *p <* 0.0001, 374% increase) and BB (*t*_14_ *=* 3.2, *p =* 0.006, 58% increase) muscle significantly increased their activity relative to baseline. For wrist extension movements (Figure 7F), a one-sample t-test showed that both the WE (*t*_14_ = 5.3, *p* = 0.0001, 280% increase) and TB (*t*_14_ = 3.5, *p* = 0.003, 72% increase) muscle significantly increased their activity relative to baseline.

**Figure 7:**
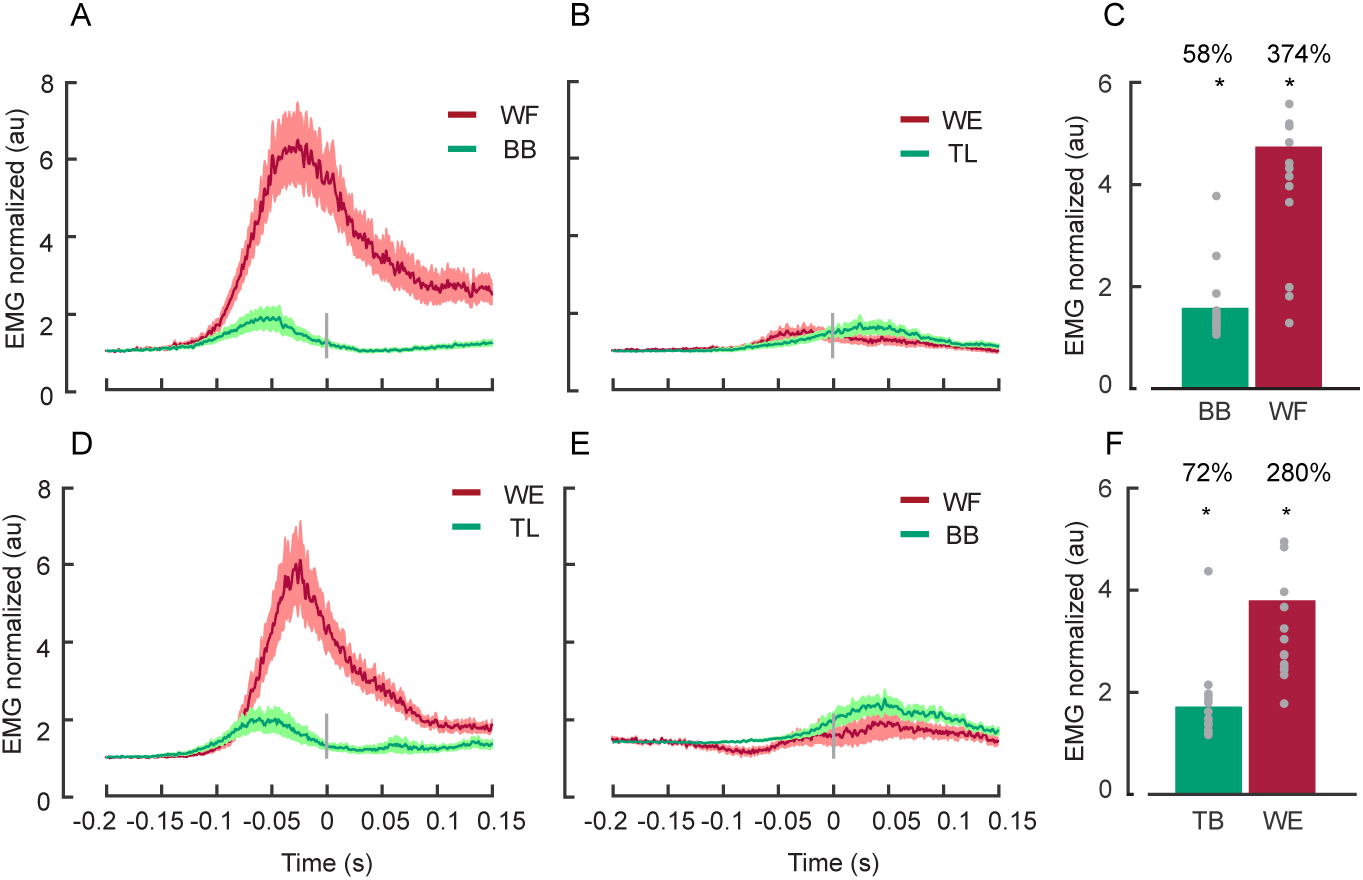
Muscle Activity, Experiment 2. (A) Normalized mean muscle activity of BB and WF muscles and (B) Normalized mean muscle activity of their respective antagonist muscles (TB and WE muscles) during a wrist flexion movement. Data are aligned on movement onset. Shaded areas represent the standard error of the mean. (C) Mean agonist muscle activity (-100 to 100ms, relative to movement onset) is shown. (D-F) Extension movements are shown using the same layout as (A-C).

As evident from the average muscle responses in Figure 7, we found that both BB (*t*_14_ = –33.7, *p* < 0.0001) and WF (*t*_14_ = –26.2, *p* < 0.0001) muscle activity significantly preceded wrist flexion movement onset by 100 ms (SE 3), and 90 ms (SE 3), respectively (Figure 8A). Similarly, we found that TB (*t*_14_ *=* –21.8, *p* < 0.0001) and WE (*t*_14_ *=* –16.3, *p* < 0.0001) muscle activity significantly preceded wrist extension movement onset by 100 ms (SE 4), and 70ms (SE 4), respectively (Figure 8C). We further investigated whether there was a difference in onset times between muscles using paired t-tests (one for wrist flexion and one for wrist extension movements) applied to average onset times. We found a reliable difference in onset times for both wrist flexion and extension movements (flexion: *t*_14_ = –2.3, *p* = 0.003; extension: *t*_14_ = –5.9, *p* < 0.0001) whereby elbow muscle activity preceded the wrist muscle activity (Figure 8 B,D).

**Figure 8:**
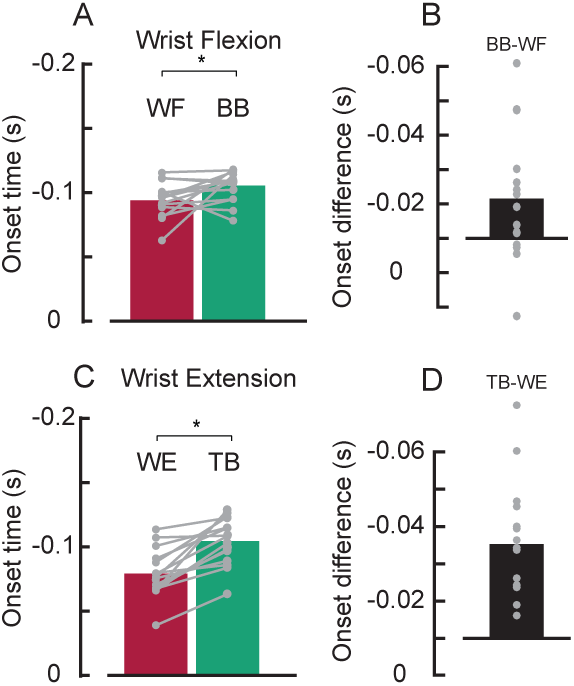
Muscle Onset Timing, Experiment 2. (A) Mean onset times of BB and WF muscles during flexion movements are shown for Experiment 2. Gray lines indicate data from individual participants. Asterisks indicate reliable effects (*p <* 0.05, see main text). (B) Difference of onset times between BB and WF muscles are shown. Dots represent data from individual participants. (C,D) Wrist extension movements are shown using the same format as (A,B).

Lastly, we examined the correlation between elbow and wrist muscle activity on a trial-by-trial basis. As in Experiment 1, we expected a positive relationship between elbow and wrist muscle activity in terms of both amplitude and onset timing. A one-sample Wilcoxon Ranksum on individual correlation coefficients revealed a reliable positive relationship between elbow and wrist muscle activity magnitude for wrist flexion trials (*W*_14_ = 116, *p <* 0.0004; 9 participants showed individually significant positive correlations) but not wrist extension movements (*W*_14_ = 77, *p* = 0.35; 5 participants). We also found reliable positive relationships between elbow and wrist onset timings for both wrist flexion and extension movements (flexion: *W*_14_ = 120, *p* < 0.0001; extension: *W*_14_ = 120, *p* < 0.0001; for both flexion and extension, all participants showed significant positive correlations).

### Experiment 3: Compensating for interaction torques introduced by hand orientation

We assessed whether shoulder muscle activity compensates for differences in the interaction torques introduced by changing hand orientation during single-joint elbow movements. That is, participants made the same elbow movement as in Experiment 1 with their wrist beginning in three different configurations and we tested whether shoulder muscle activation accounted for how hand orientation influenced interaction torques at the shoulder caused by forearm rotation. As expected, participants completed the task mostly via single-joint rotations at the elbow joint (Figure 9) and they had little difficulty reaching the goal target within the imposed speed and accuracy constraints (mean success rate = 95%).

**Figure 9:**
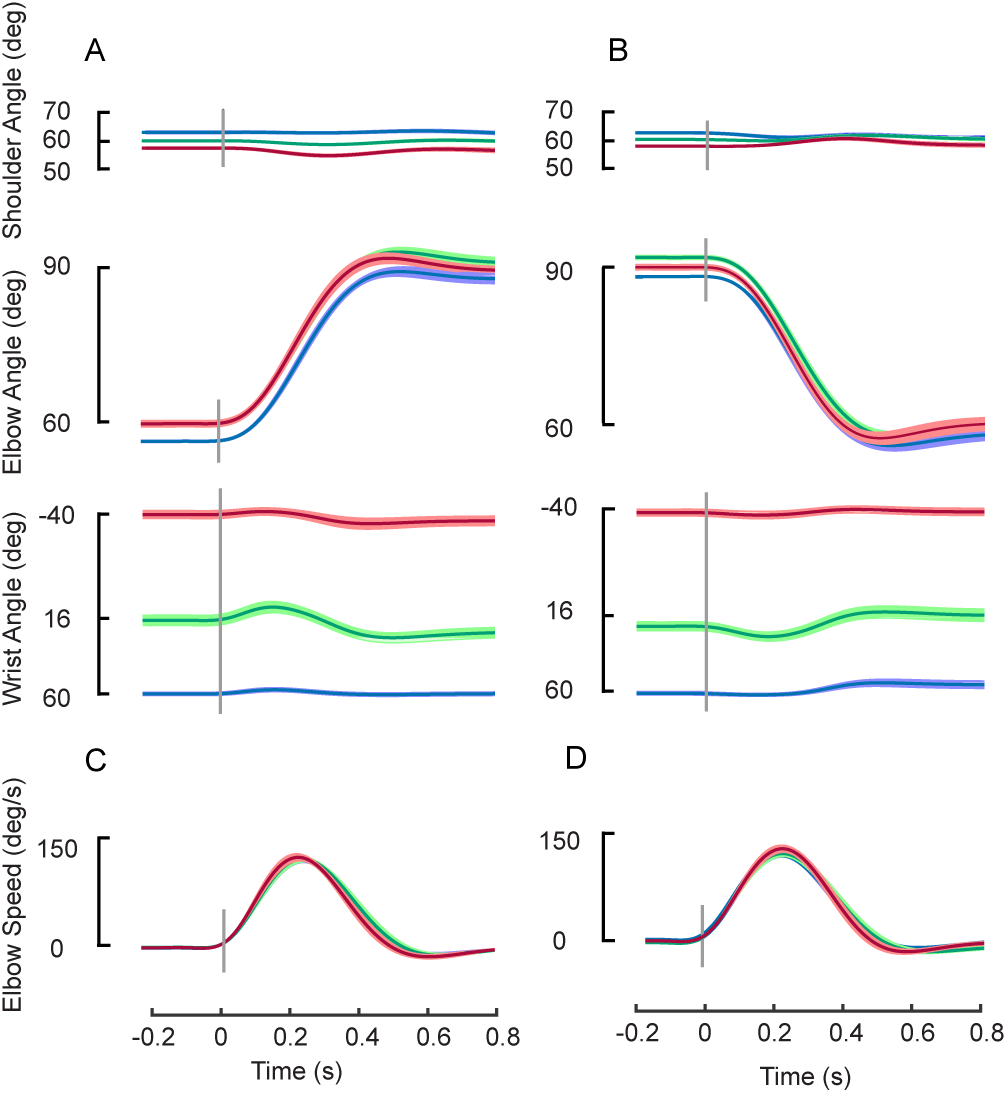
Reaching Behavior, Experiment 3. (A) Kinematics of the shoulder, elbow and wrist joints and (C) Velocity profiles of the elbow joint during flexion movements in Experiment 3. Data are aligned on movement onset. Shaded areas represent the standard error of the mean. (B and D) Same as (A and B) but for elbow extension movements, respectively.

Figure 10 illustrates the average agonist muscle activity at the PEC and PD muscles during elbow flexion (A) and extension (C), respectively. Qualitatively, shoulder muscle activity scaled for the three different initial wrist configurations and this scaling was consistent with the magnitude of interactions torques introduced by changing hand orientation. We quantified this relationship by calculating the magnitude of shoulder muscle activity in each hand orientation condition in a fixed time window (-200 to 100ms) relative to movement onset. We performed linear regression for each muscle sample to determine whether there was a reliable relationship between shoulder muscle activity and the initial wrist configurations. A one-sample t-test of the individual slopes revealed a reliable negative slope for PEC muscle activity during elbow flexion (*t*_14_ = –5.9, *p <* 0.0001; Figure 10B) and PD muscle activity during elbow extension (*t*_14_ = –5.0, *p* = 0.0001; Figure 10D).

**Figure 10:**
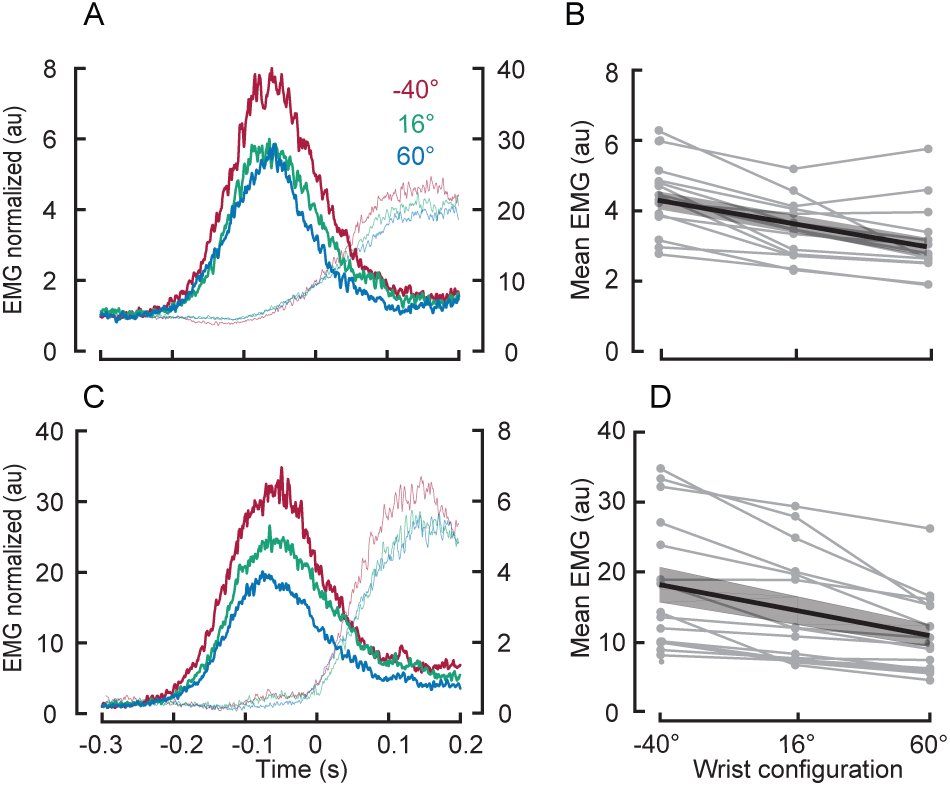
Muscle Activity, Experiment 3. Shoulder muscles compensate for the magnitude of interaction torques introduced by changing wrist configuration. (A) Average PEC muscle activity is shown during elbow flexion movements in three different wrist configurations. Data are aligned on movement onset. (B) Individual data (gray lines) and mean regression slope (black line; shaded area = SE) between mean agonist muscle activity (-200 ms to 100 ms relative to movement onset) and the three wrist configurations. (C and D) Same format as (A and B) but for elbow extension movements, respectively.

Figure 11A and B show the cumulative distribution of onset times of the relevant shoulder, elbow and wrist muscles for elbow flexion and extension movements, respectively. We investigated differences in onset times across muscles and, in this case, hand orientation with a two-way repeated measures ANOVAs (one for elbow flexion and one for elbow extension). For elbow flexion movements, we found a reliable effect of muscle (*F*_2,117_ = 5.4, *p* = 0.005), but no effect orientation (*F*_2_,_117_ = 0.25, *p* = 0.78) and no interaction (*F*_4,117_ = 0.055, *p* = 0.99). Tukey post-hoc tests showed that PEC muscle activity reliably preceded BB muscle activity (*p* = 0.001) by 20 ms (*p* = 0.0011) and that BB muscle activity preceded WF muscle activity by 7 ms (*p* = 0.0016). For elbow extension movements, we found a reliable effect of orientation (*F*_2,117_ = 4.0, *p* = 0.02) as well as muscle (*F*_2,117_ = 5.2, *p* = 0.006) but no interaction (*F*_4,117_ = 0.96, *p* = 0.42). Tukey post-hoc tests showed that muscle onset times in the −40°hand orientation preceded those in the 16°hand orientation by 25ms (*p* = 0.0026) and the 60°hand orientation by 24ms (*p* = 0.0053). In addition, Tukey post-hoc tests showed that TB muscle activity preceded PD muscle activity by 9 ms (*p* = 0.0007), and that TB muscle activity preceded the WE muscle activity by 22 ms (*p* = 0.005).

**Figure 11:**
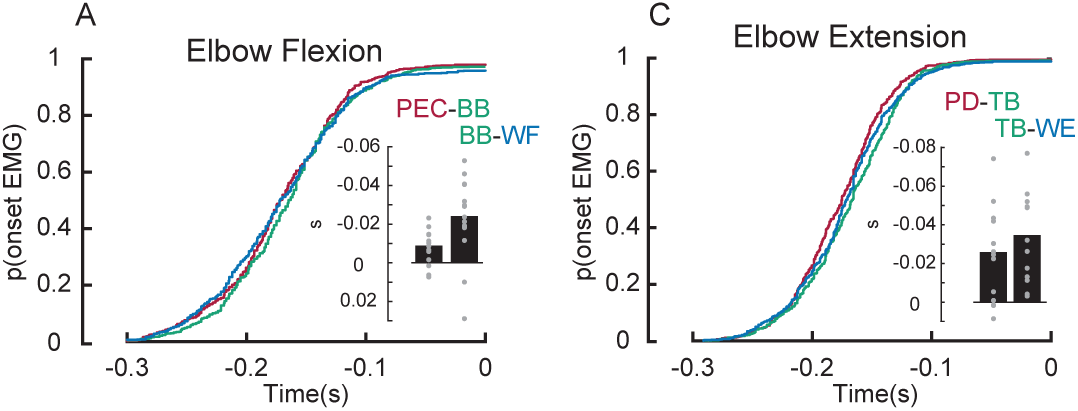
Muscle Onset Timing, Experiment 3. (A). Cumulative distributions of onset times of shoulder, elbow and wrist muscles during flexion movements are shown for Experiment 3. Insets represent the group average onset timing difference between muscle pairs. Each gray dot represents mean onset time differences from a single participant. (B) Same as (A) but for elbow extension movements.

### Experiment 4: Rapid feedback responses at the shoulder account for interaction torques caused by hand orientation

Our last experiment mimicked Experiment 3 but we examined feedback control. That is, we tested whether rapid feedback responses to mechanical perturbations compensate for interaction torques arising due to the orientation of the hand. We did so by extending our previously established paradigm (Kurtzer et al., 2008) to the three-joint situation. Briefly, for two wrist configurations we applied shoulder and elbow torques that yielded very similar shoulder and elbow motion (see Methods). This allowed us to directly test whether rapid feedback responses in shoulder muscles modulated their muscle activity according to the underlying torque at the shoulder, which differs because of the intersegmental effects introduced by the different wrist configurations.

Figure 12 shows the average kinematics for each participant at the shoulder, elbow and wrist joints with the wrist initially in flexion (60°, A) or extension (-40°, B). Note the highly similar motion at the shoulder (median difference = 0.003°) and elbow (0.04°) for the two hand orientations, which was achieved by applying higher shoulder torque for the wrist extension condition (see Methods, Figure 12C,D).

**Figure 12:**
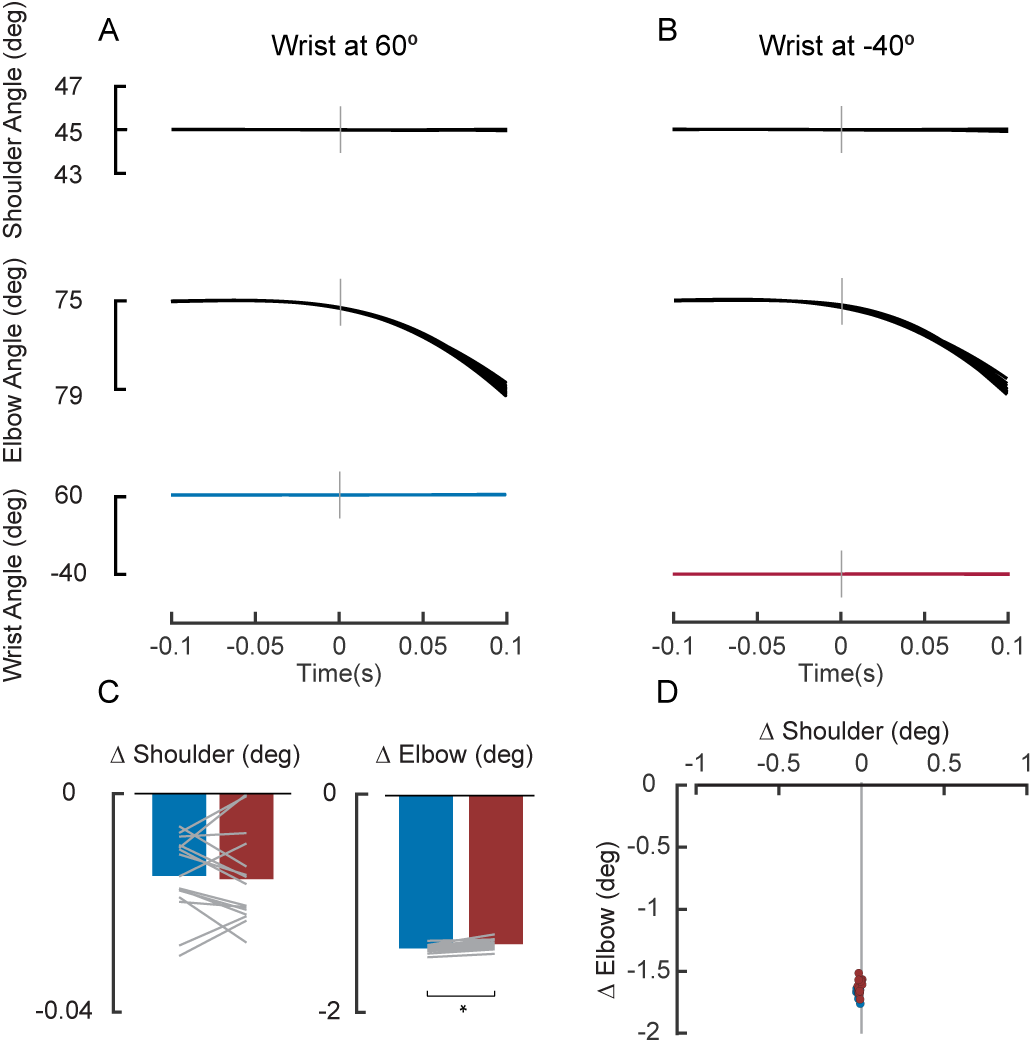
Perturbation evoked movement, Experiment 4. (A) Kinematics of the shoulder, elbow and wrist joints are shown following a multi-joint perturbation when the wrist was positioned at 60°. (B) Same layout as (A) but when the wrist was positioned at −40°. (C) Mean shoulder and elbow joint displacement at 50 ms post perturbation is shown for Load Combination 1 (see Methods). Gray lines indicate data from individual participants. (D) Shoulder and elbow joint angles at 50 ms post-perturbation are shown for Load Combination 1.

Figure 13A presents the average shoulder (PEC) muscle activity for the two initial hand orientations. We focused on the shoulder as the elbow had the same underlying torques for both hand orientations. Qualitatively, we observed no evoked shoulder activity during the short-latency epoch. This is expected given that the perturbation caused almost no local shoulder motion. In addition, the mean response in the long latency and voluntary epochs showed larger responses for the wrist configuration that involved greater underlying shoulder torque. Statistical analysis confirmed that this was a highly reliable pattern. Paired t-tests showed no significant difference in average muscle activity as a function of load condition within the short latency epoch (*t*_14_ = –0.5*,p =* 0.626, Cohen’s *d =* 0.128) but did reveal reliable effects in the long-latency (*t*_14_ = –4.1, *p* = 0.001, *d* = 1.05) and voluntary epochs (*t*_14_ = –3.7, *p* = 0.002, *d* = 0.95) that were appropriate for countering the applied shoulder torques (Figure 13B).

**Figure 13:**
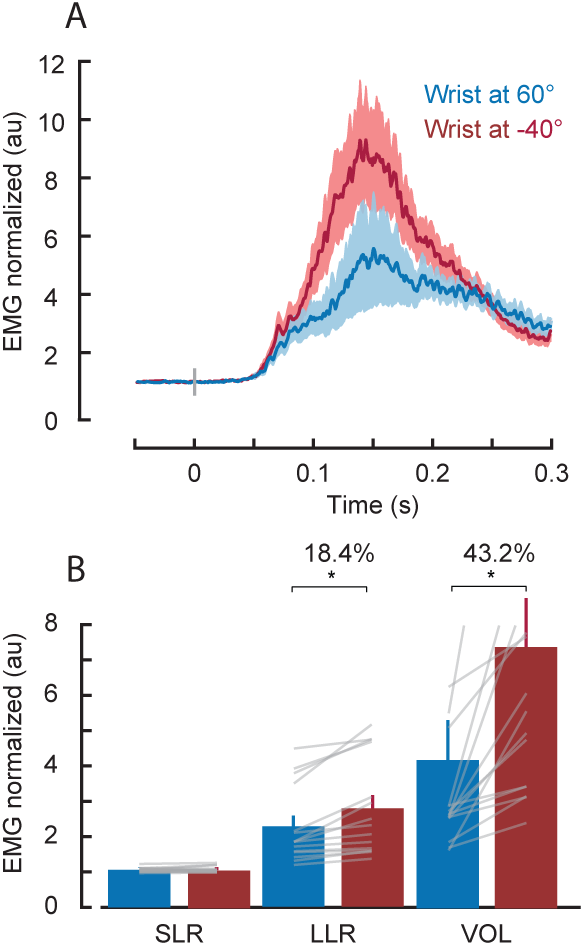
Fast Feedback Responses, Experiment 4. (A) Mean PEC muscle activity is shown for two different hand orientations, for Load Combination 1 (See Methods). Data are aligned on perturbation onset. Shaded areas represent the standard error of the mean. (B) Bars represent the mean muscle activity in pre-defined epochs relative to perturbation onset: short latency (0-50 ms), long latency (50-100 ms) and voluntary (100-150 ms) responses. Gray lines represent individual participants. Asterisks depict reliable effects (*p <* 0.05, see Main Text).

## DISCUSSION

### Summary

We examined how robustly the nervous system accounts for intersegmental dynamics across the shoulder, elbow and wrist joints during both feedforward (i.e. self-initiated) and feedback (i.e. reflexive) control. In Experiment 1, we found that shoulder muscle activation predictively scaled according the magnitude of upcoming interaction torques, both as a function of elbow configuration and movement speed. In Experiment 2, we found that elbow muscle activation predictively scaled to compensate for interactions torques during single-joint wrist movement. In Experiment 3, we found that shoulder muscle activity predictively scaled according to the interaction torques introduced by different hand orientations during single-joint elbow movement. In Experiment 4, we found that feedback responses at the shoulder evoked by a mechanical perturbation that caused single-joint elbow motion also accounted for hand orientation, starting approximately 50 ms after perturbation onset (i.e. within the long-latency epoch). Taken together, our results demonstrate that the nervous system robustly accounts for intersegmental dynamics across the proximal to distal musculature of the arm, and does so for both feedforward and feedback control.

### Accounting for interaction torques during feedforward control

Many studies have investigated how the nervous system deals with interaction torques during self-initiated reaching by having participants make single joint movements when multiple joints are free to move (Almeida et al., 1995; Corcos et al., 1989; Galloway and Koshland, 2002; Gottlieb, 1998; Gribble and Ostry, 1999; Koshland et al., 1991). These studies have established that muscles spanning joints that are adjacent to the moving joint contract prior to movement onset and, as such, predictively compensate for the interaction torques about to arise because of the movement. Such compensation has been demonstrated for shoulder muscles during pure elbow movements (Almeida et al., 1995; Gribble and Ostry, 1999; Corcos et al., 1989; Galloway and Koshland, 2002), elbow muscles during pure shoulder movements (Almeida et al., 1995; Gribble and Ostry, 1999; Galloway and Koshland, 2002), and wrist muscles during pure elbow movements (Koshland et al., 1991). The results of our first two experiments confirm and extend these previous findings by demonstrating that muscle activity at the shoulder joint scale according to the predicted magnitude of the interaction torques during single joint elbow movement in different elbow configurations and movement speeds (Experiment 1), and muscle activity at the elbow joint scales according to interaction torques introduced by single joint wrist movement (Experiment 2).

Accounting for intersegmental dynamics becomes more complex in a three-joint scenario because interaction torques arise at all joint segments. We investigated whether the nervous system accounts for interaction torques across three joints by having participants perform single joint elbow movements with different wrist configurations (Experiment 3). Doing so requires generating different torques at the shoulder as a function of the hand orientation. An important point to note is that these torque demands are substantially smaller than the torque demands created by the two-joint situations studied in Experiments 1 and 2. Thus, it seemed possible the nervous system would use a qualitatively different control strategy to counter these smaller torque demands. For example, the nervous system could have countered these interaction torques by co-contracting shoulder agonist and antagonist muscles, thereby increasing the stiffness of the joint and limiting shoulder motion. Indeed, the control of limb stiffness can be an important control scheme in unstable environments (Burdet et al., 2001; Franklin et al., 2007; Hogan, 1985; McIntyre et al., 1996; Milner, 2004). However, we found little evidence of co-contraction. Rather, we observed marked phasic activity of shoulder agonist muscles that predicted the magnitude of interaction torques introduced by wrist configuration (Figure 10). These findings suggest that accounting for intersegmental dynamics is a core computation for the feedforward control of reaching movements that is evident for even very small changes in speed and limb configuration across multiple joints.

### Predicting interaction torques during feedforward control

Previous findings have emphasized that during single joint movement, muscles at the stationary joint are activated prior to movement onset, indicating that the nervous system compensates for interaction torques in a predictive manner (Almeida et al., 1995; Gribble and Ostry, 1999; Koshland et al., 1991; Sainburg et al., 1995, 1999). Our findings are consistent with this type of organization (Figure 4, 8, 11). More interestingly, Gribble and Ostry (1999) demonstrated a particular temporal ordering of muscle activation for single joint movements whereby proximal muscles were activated prior to distal muscles. Specifically, they found that the onset of shoulder muscle activity preceded not just elbow movement but also the onset of elbow muscle activity by 20-50 ms, a finding also evident at the level of single neurons in the monkey primary motor cortex (Fetz et al., 1989; Humphrey, 1972; Scott, 1997). Although our results are broadly consistent with this proximal to distal rule, the timing differences we found were either substantially smaller than previously reported (<20 ms) and/or did not reach statistical significance (Figure 4, 8).

One possibility is that the proximal to distal differences in recruitment timing merely reflect the conduction delays associated with neural commands propagating to more distal muscles. This simple explanation is attractive but seems unlikely given that such delays would account for substantially less than ~5ms (Ingram et al., 1987; Wang et al., 1999). Moreover, it should be emphasized that our onset time estimates (and most previous studies showing similar effects) may be biased in a way that under-estimates onset timing differences between distal and proximal muscles. That is, techniques that estimate muscle onset time as when the EMG signal reaches some arbitrary threshold relative to baseline will be biased towards earlier values for muscles showing greater activation levels - the more distal muscles in our experiments. We thus consider our estimate of onset timing differences a lower bound on how much the onset of activity in the distal muscles leads proximal muscles in our task.

It is important to note that the proximal-distal rule is not mandatory. For example, a large body of work in object manipulation has shown that grip forces used to stabilize hand held objects are modulated roughly in phase with the self-generated forces arising during arm movements with object manipulation (Flanagan and Wing, 1997; Danion and Sarlegna, 2007; Diamond et al., 2015; Hadjiosif and Smith, 2015; Wolpert and Flanagan, 2001). So why are proximal muscles activated prior to distal muscles in the simple reaching task used in the present experiments? Previous work has suggested that the multi-joint movement is organized in a hierarchical structure with leading and subordinate joints (Dounskaia, 2005). Muscles at the leading joint act to accelerate the limb as during single joint movement and muscles at the subordinate joint act to regulate interaction torques and tune net torque to create the desired movement. This simplifying control structure is appealing in many respects but does not provide a clear explanation for our findings. That is, if the distal joint is the leading joint, which seems sensible given that this is the only segment with net acceleration, then according to the leading joint hypothesis it should be activated prior to the proximal joint that needs to regulate interaction torques to keep the hand on target. Gribble and Ostry (1999) proposed the more general idea that a proximal to distal rule might reflect an organizational strategy for maintaining limb stability. However, from a purely mechanical perspective there is no clear reason to pre-activate proximal muscles to counter interaction torques arising from distal joint rotation. One possibility is that onset differences could reflect different force recruitment properties of muscles spanning these different joints. Indeed, previous work in monkeys reveals a relatively high abundance of fast twitch muscle fibers in superficial elbow flexors (Singh et al., 2002) and it is the flexor condition where we see the largest and most reliable lead in shoulder muscle onset times. Although this explanation is appealing for the shoulder and elbow, it does not necessarily support a general proximal-distal rule for arm muscles. Additional work is clearly needed to establish the general validity of the proximal-distal rule and to link it to a specific mechanism.

### Accounting for interaction torques during feedback control

Many studies have shown that the nervous system accounts for the limb’s intersegmental dynamics during self-initiated (i.e. feedforward) control of the arm (Almeida et al., 1995; Cooke and Virji-Babul, 1995; Corcos et al., 1989; Galloway and Koshland, 2002; Gottlieb, 1998; Gribble and Ostry, 1999; Gritsenko et al., 2011; Hollerbach and Flash, 1982; Koshland et al., 1991; Pigeon et al., 2013; Sainburg et al., 1995, 1999; Virji-Babul and Cooke, 1995). A related series of studies has demonstrated that this capacity is also present during reflexive (i.e. feedback) control of the arm (Crevecoeur et al., 2012; Kurtzer et al., 2008, 2009, 2014, 2016; Lacquaniti and Soechting, 1984, 1986a, b; Pruszynski et al., 2011; Soechting and Lacquaniti, 1988). For example, Soechting and Lacquaniti (1988) investigated rapid feedback responses following mechanical perturbations at the shoulder and elbow joints when both joints were free to move. They showed that feedback responses starting approximately 20 ms after perturbation onset (i.e. the short-latency stretch response) reflected motion at one joint whereas those starting approximately 50 ms after perturbation onset (i.e. the long-latency stretch response) reflected motion at multiple joints. Kurtzer and colleagues 2008 further investigated this issue by applying a specific combination of mechanical perturbations at the shoulder and elbow joints that led to either (i) the same motion at the shoulder and different motion patterns at the elbow or (ii) minimal motion at the shoulder and different motion patterns at the elbow. Their results showed that the short-latency stretch response responded only to local joint motion whereas the long-latency stretch response accounts for the limb’s intersegmental dynamics and responds to the underlying applied torques.

Our present work (Experiment 4) tested whether feedback control of the arm accounts for the limb’s intersegmental dynamics across the whole arm. We did so by applying two sets of loads to the shoulder and elbow that yielded similar motion profiles at the shoulder and elbow joints when the wrist was locked in two different configurations. As in Experiment 3, the applied loads were quite similar because the mass of the hand has a subtle influence on the overall mechanical properties of the arm. Nevertheless, we saw robust changes during the long-latency stretch response that were appropriate for countering the underlying applied loads with the wrist positioned in different configurations. Thus, it appears that the nervous system is highly sensitive to intersegmental dynamics across the shoulder, elbow and wrist joints during both feedforward and feedback control.

The functional similarity between feedforward and feedback control in our study lends further support to the idea that these responses engage a similar neural circuit. One likely node in this circuit is primary motor cortex. Gritsenko and colleagues 2011 used transcranial magnetic stimulation over the human primary motor cortex during self-initiated reaching movements towards targets that included assistive or resistive interaction torques. Their results showed that motor evoked potentials were greater for movement directions that included resistive interaction torques as compared to assistive movements, suggesting that M1 indeed mediates feedforward compensation for the limb’s intersegmental dynamics. Similarly, Pruszynski and colleagues 2011 showed that transcranial magnetic stimulation over M1 in humans can potentiate shoulder muscle responses after mechanical perturbations that cause pure elbow displacement, suggesting that M1 mediates feedback compensation for the limb’s intersegmental dynamics. An important area of future research, which we are actively pursuing, is determining whether the same neurons in M1 carry these signals for feedforward and feedback control, and whether these signals reflect processing intrinsic to M1 or whether they reflect computations performed in other parts of the brain.

